# SARS-CoV-2 promotes microglial synapse elimination in human brain organoids

**DOI:** 10.1101/2021.07.07.451463

**Authors:** Samudyata, Ana Osório Oliveira, Susmita Malwade, Nuno Rufino de Sousa, Sravan K Goparaju, Jessica Gracias Lekander, Funda Orhan, Laura Steponaviciute, Martin Schalling, Steven D Sheridan, Roy H. Perlis, Antonio Rothfuchs, Carl M. Sellgren

**Affiliations:** Department of Physiology and Pharmacology, Karolinska Institute, Stockholm, Sweden; Department of Microbiology, Tumor and Cell Biology, Karolinska Institute, Stockholm, Sweden; Department of Molecular Medicine and Surgery, Karolinska lnstitutet and Center for Molecular Medicine, Karolinska University Hospital, Stockholm, Sweden; Center for Genomic Medicine and Department of Psychiatry, Massachusetts General Hospital, Boston, MA, USA; Centre for Psychiatry Research, Department of Clinical Neuroscience, Karolinska lnstitutet & Stockholm Health Care Services, Stockholm County Council, Karolinska University Hospital, Stockholm, Sweden

## Abstract

Neuropsychiatric manifestations are common in both the acute and post-acute phase of SARS-CoV-2 infection, but the mechanisms of these effects are unknown. In a newly established brain organoid model with innately developing microglia, we demonstrate that SARS-CoV-2 infection causes an extensive cell death and loss of post-synaptic termini. Despite limited neurotropism and a decelerating viral replication, we observe a threefold increase in microglial engulfment of postsynaptic termini after SARS-CoV-2 exposure. We define the microglial responses to SARS-CoV-2 infection by single cell transcriptomic profiling and observe an upregulation of interferon-responsive genes as well as genes promoting migration and synaptic stripping. To a large extent, SARS-CoV-2 exposed microglia display a transcriptomic profile previously observed in neurodegenerative disorders characterized by early a synapse loss and an increased incident risk after a Covid-19 infection. Our results reveal that brain organoids infected with SARS-CoV-2 display disruption in circuit integrity via microglia-mediated synapse elimination and identifies a potential novel mechanism contributing to cognitive impairments in patients recovering from Covid-19.

## INTRODUCTION

A significant proportion of patients infected with severe acute respiratory syndrome coronavirus 2 (SARS-CoV-2), the causative agent of coronavirus disease 2019 {COVID-19), display acute neurological and psychiatric symptoms^1^. Despite a low neural expression of the major receptor (angiotensin-converting enzyme 2; ACE2) for SARS-CoV-2 entry, autopsy studies have also detected the presence of SARS-CoV-2 RNA and protein in brain tissue of around half of the examined COVID-19 patients^2^. Survivors then commonly report persistent central nervous system (CNS)-related symptoms, such as memory loss and sleep disturbances^3^, as well as depressive symptoms^4^. In line with these findings, a subset of the patients also demonstrate widespread disruptions to micro-structural and functional brain integrity in the recovery stages^5^. Further, the risk of receiving a neurological or psychiatric diagnosis related to cognitive impairment or behavioral abnormalities, including dementia, parkinsonism, or a psychotic disorder, is increased also in non-hospitalized patients up to at least 6 months after infection, as compared to other respiratory tract infections and with influenza^6,7,3^. Long-lasting neurobehavioral and cognitive impairments are also common features of other RNA viruses with a neuro-invasive potential, with the CNS-damaging processes also continuing after virus elimination^8^. So far, the mechanisms causing neurocognitive sequelae after viral encephalitis are poorly understood, but recent studies in rodents suggest that interferon-responses in microglia can lead to excessive microglial synapse elimination and disruption of neuronal circuit integrity^9,10^. To some extent, this could explain why the developmental period of infection influences the extent of neurocognitive sequelae^11^, as microglial functions involving circuit refinement exhibit a clear temporal pattern during neurodevelopment^12^.

Human brain organoids have been successfully employed to study neurotropism and neurotoxic effects of viruses such as the Zika virus^12,13^, and now more recently SARS-CoV-2^14–18^. However, as microglia are of non-ectodermal origin^19,20^, and the predominate germ layer fate in embryoid bodies is ectodermal, brain organoids typically lack innately developing microglia. This has limited the mechanistic understanding of microglial cellular responses in viral CNS infection, such as in COVID-19, as well as understanding of cellular responses of astrocytes and neurons in the context of surveying microglia.

In this study, we address this limitation by using a newly established protocol for generating undirected brain organoids with innately developing microglia that arise from Brachychury^+^ mesodermal progenitors^21^. Despite a low multiplicity of infection (MOI) resulting in a modest infection, we observe an extensive neuronal cell death and a decrease in postsynaptic density secondary to microglial synaptic stripping. We perform single cell RNA sequencing (scRNA-seq) to characterize the temporal dynamics of single cell responses in infected organoids, and observe a cluster of interferon responsive microglia, also defined by an upregulation of genes promoting migration and microglial synapse elimination, that largely overlap with the transcriptomic profile observed in neurodegenerative disorders characterized by an early synapse loss and an increased incident risk after a Covid-19 infection. These results indicate that SARS-CoV-2 has the potential to disrupt neuronal circuit integrity via microglia-mediated synapse elimination and identifies a potential novel mechanism contributing to cognitive impairments in patients recovering from Covid-19.

## RESULTS

### Limited neurotropism but extensive cell death and synapse loss in infected brain organoids

With a few modifications (see **Methods** and **Fig. la**), the protocol reported by Ormel et al.^21^ was adapted to miniaturized bioreactors and used to generate undirected brain organoids. After 56 days in vitro (DIV), these undirected organoids were found to contain neural progenitor cells (NPCs; PAX6^+^), neurons (β-tubulin^+^ and MAP2^+^), neural crest cells (SOX10^+^), cells of astrocytic lineage (GFAP^+^ and S100B^+^ at 56 DIV and AQP4^+^ at 110 DIV; **Supplementary Fig. la**), along with microglia (IBA1^+^ and CD68^+^); **Fig. lb.** We exposed such organoids to active SARS-CoV-2 virus for 2h at an estimated MOI of 0.3, after which they were washed and transferred into fresh media to monitor the course of infection. Supernatants were harvested at 6, 24, 48, 72 hours post-infection (hpi) for viral replication assessment. Viral RNA-dependent RNA polymerase *(Rdrp)* and nucleocapsid (N) transcription were measured in supernatants via RT-qPCR and we observed a time dependent increase for both these transcripts **(Fig. 2a** and **Supplementary Fig. lb**), indicating the virus is capable of initiating RNA replication of its genomic material within the cells of the organoid, but at a slower pace as compared to Vero-E6 cells **(Supplementary Fig. lb).** To validate SARS-CoV-2 could assemble and release complete viral particles in the organoids, we performed plaque formation assays using supernatants from infected organoids and observed an release of infectious viral particles as early as 6hpi, although the number of plaque forming units (PFUs) then quickly decreased **(Fig. 2b).**

**Figure 1:**
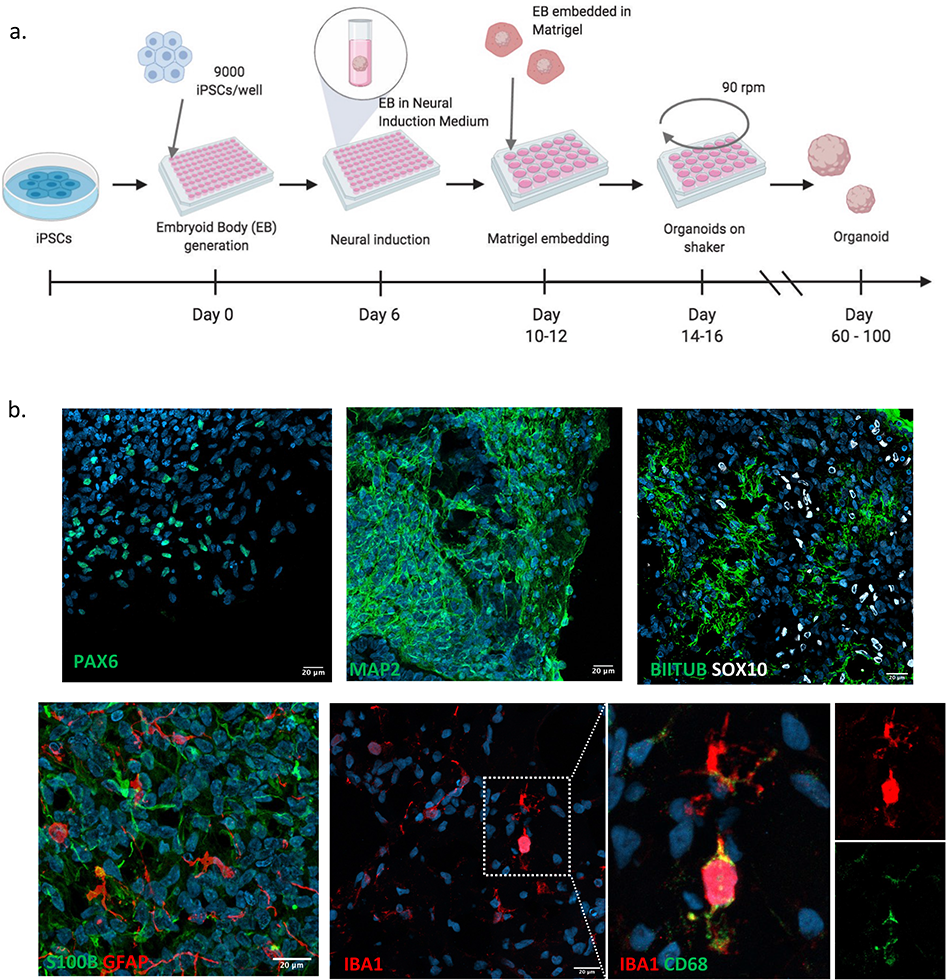
Cellular composition of DIV 56 brain organoids containing innately developing microglia. (a) Schematic of brain organoid generation using an undirected protocol adapted from Ormel et al., (b) IHC staining of different cell type markers showing the presence of NPCs (PAX6), immature and mature neurons (B-111 Tubulin and MAP2), schwann cells (SOXlO), astrocytic lineage cells (GFAP, Sl00B) and microglia (IBAl, CD68).

**Figure 2:**
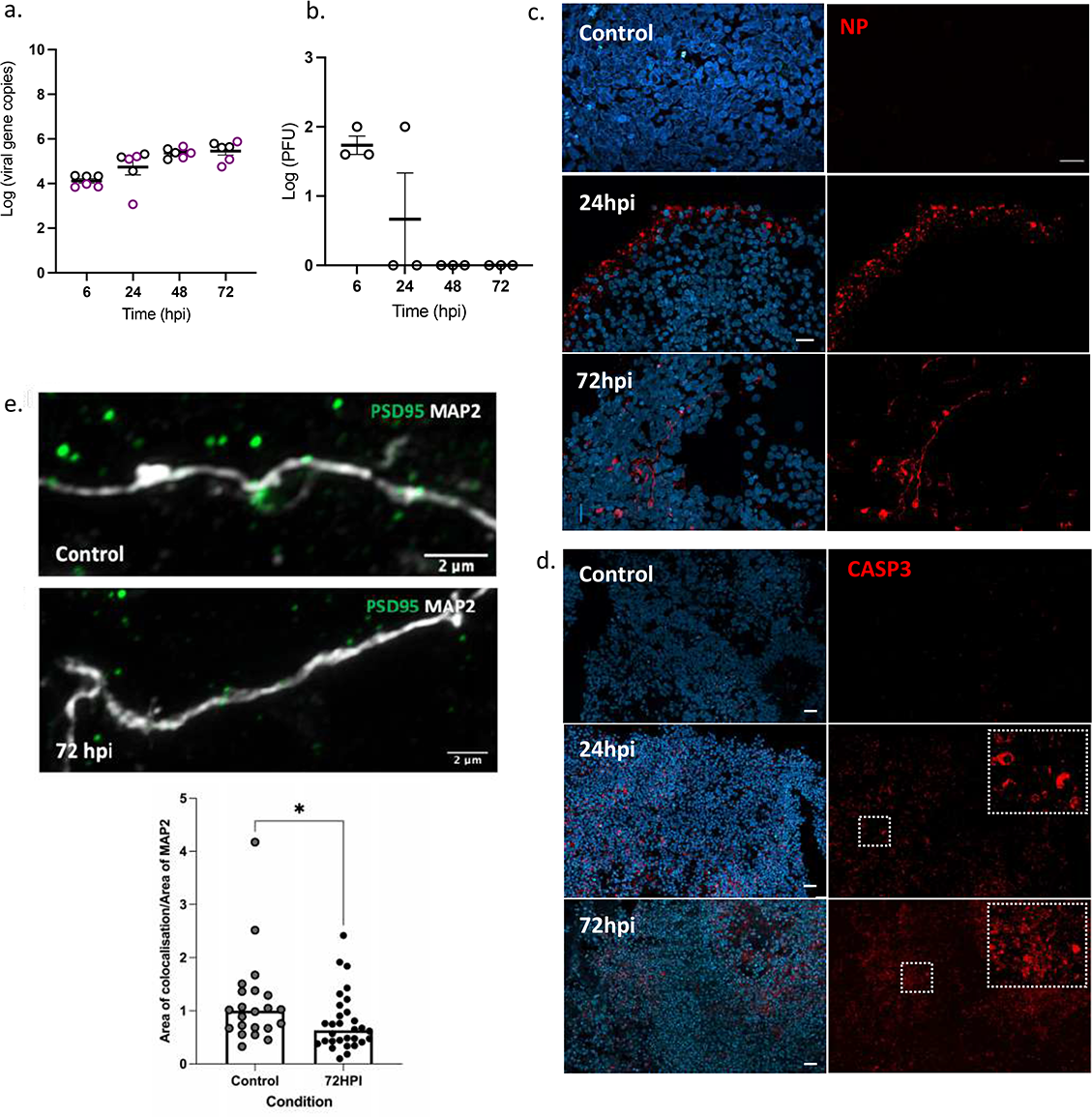
SARS-CoV-2 infection of immunocompetent brain organoids. Brain organoids infected with live SARS-CoV-2 virus was collected at different time points following exposure. (a) qPCR analyses on supernatants of infected brain organoids at 56 DIV (n=3, black circle) and 110 DIV (n=3, magenta circle) showing log fold change in viral nucleocapsid *(N)* gene copies, at different time points. Error bars indicate standard errors of the mean (SEM). **(b)** Log transformed PFUs of supernatants from infected brain organoids at different time points. Error bars indicate SEM. Representative IHC tiled images showing staining of **(c)** nucleocapsid protein (NP) and **(d)** cleaved caspase 3 (CASP3) in 56 DIV infected brain organoids and uninfected control organoids. **(e)** Postsynaptic density (PSD-95) decreased by 50% in organoids at 72hpi (Mann-Whitney U-test: P=0.025). Images are representative for each condition. In the graph, bars display medians, and the data is normalized to median uptake in control organoids (n=3 organoids per condition). Each data point represents the mean of all cells per field of view.

At two time points (24 and 72hpi), we fixed infected and uninfected organoids (56 DIV) for immunohistochemical (IHC) analyses. At 24hpi, a strong signal for the viral nucleocapsid protein (NP) was observed mainly in the periphery of the infected organoids, whereas at 72hpi, we observed NP staining in cellular cytoplasm or processes of infected cells **(Fig. 2e).** On average, around 3% of the cells in the superficial layers of the organoids were found to be NP^+^ at 72hpi **(Supplementary Fig. le).** Cleaved caspase3 (CASP3) staining revealed extensive cell death in the infected organoids as compared to the uninfected organoids, occurring already at 24hpi **(Fig. 2d).** Casp3^+^ cells were not just limited to superficial layers like NP^+^ cells and the number of Casp3^+^ cells greatly exceeded the number of NP^+^ cells at both timepoints **(Supplementary Fig. le).** Infected organoids also displayed a clear decrease in postsynaptic density as compared to control organoids, and at 72hpi, organoids displayed a 50% decrease in post-synaptic density as measured by PSD-95 **(Fig. 2e).**

Next, we investigated infectivity and cell death in the context of cellular identity. IHC staining on infected organoids at 72hpi showed viral NP overlapping with PAX6^+^ MAP2^+^ GFAP^+^, SOX10^+^, and OLIG2^+^ cells; **Fig. 3a-f).** CASP^+^ staining was foremost observed in neurons, but also included other cell types **(Supplementary Fig. le).** In addition, we detected IBA1^+^ cells that stained positive for dsRNA **(Fig. 3f)** although the scarcity of these cells (slightly below 1% on average) obstructed meaningful quantifications. Therefore, we enriched for microglia from two infected organoids using magnetic-activated cell sorting (MACS) with CD11b beads. qRT-PCR on cell lysates suggested the capture of fetal microglia *(AIF1* expression, no *TMEM119* expression^22^ **Supplementary Fig. lf)** including viral *N* gene copies **(Fig. 3g).** Further, we derived induced human microglia-like cells (iMGs) in 2D-culture^23^, and exposed these cells to live SARS-CoV-2 virus (MOl=0.01). At 24hpi, we detected dsRNA^+^ cells **(Supplementary Fig. lg**), indicating the presence of virus inside iMGs.

**Figure 3:**
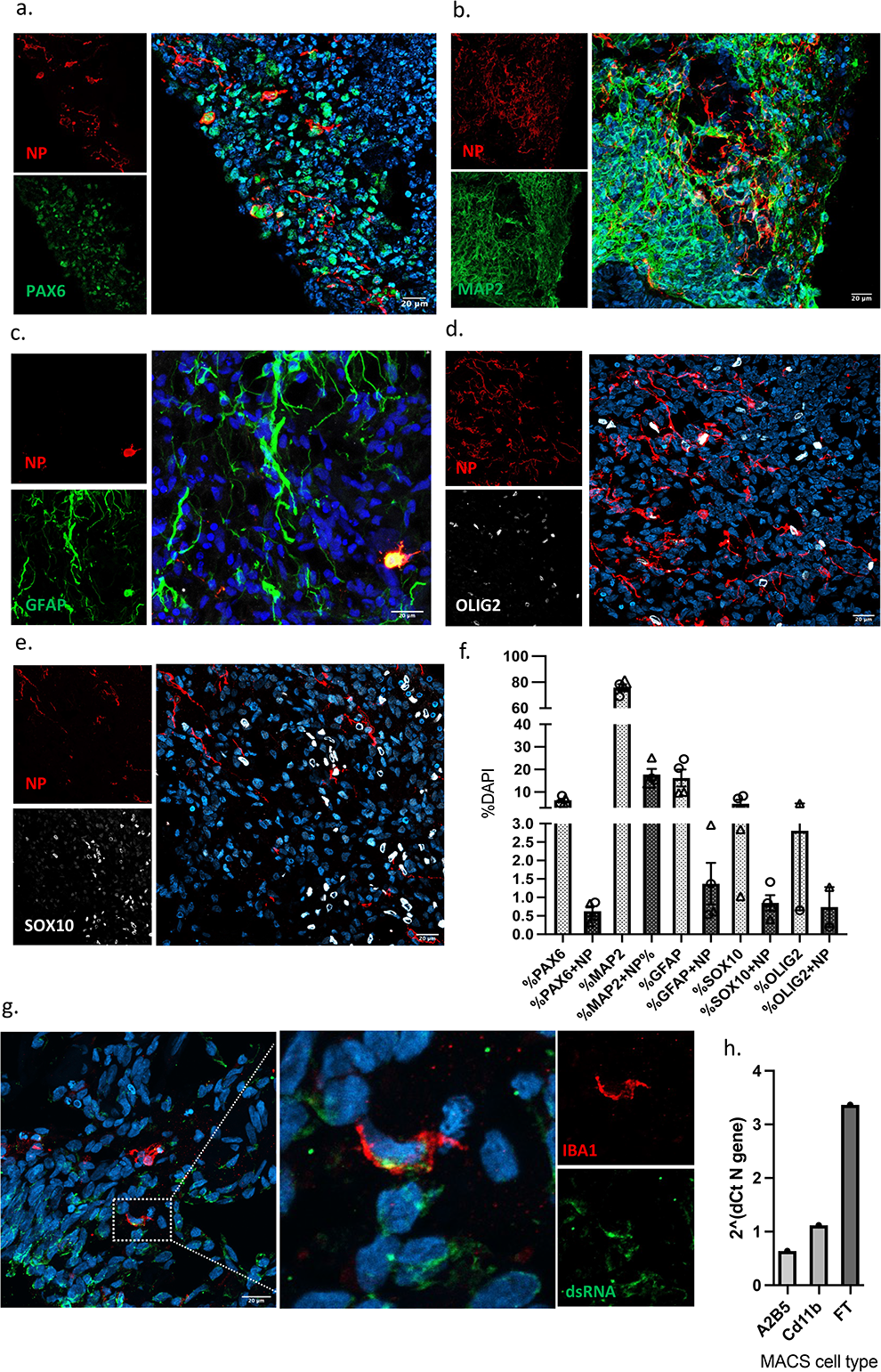
Cell type specific SARS-CoV-2 infection in brain organoids with innately developing microglia. 40X representative confocal images showing colocalization of viral NP with **(a)** neuronal precursors (PAX6), **(b)** neurons (MAP2), and a heterogenous population of progenitors marked by **(c)** GFAP **(d)** OLIG2 and (e) schwann cells (SOXlO). **(f)** Distribution of NP^+^ cells amongst different cell-type markers, expressed as percentage of DAPI from two D56 infected organoids (corresponding data points indicated by circles and triangles). Each data point is an average of 4 FOVs representing different areas on the same section. Error bars indicate SEM. (g) 40X confocal image showing colocalization of dsRNA (for viral presence) with organoid grown microglia (IBA1) and **(h)** qPCR on cell lysate fractions obtained after magnetic cell sorting (MACS) with CD11b beads, (also shown A2B5 radial glia enriched fraction) and flowthrough (FT), on infected organoids (n=2, pooled) showing presence of viral N gene copies.

### Increased microglial stripping of postsynaptic termini in infected organoids

Previous studies using murine models have shown that neuroinvasive RNA viruses can cause microglial engulfment of synaptic termini ^9,10^. To determine if SARS-CoV-2 also has the capability to induce synaptic stripping and contribute to the observed decrease in postsynaptic density observed in infected organoids, we first investigated to what extent microglia contributed to synapse remodeling in uninfected organoids (110 DIV). In line with our previous findings in 2D models^24^, super-resolution imaging revealed a baseline uptake of postsynaptic structures (indicated by PSD-95) in microglia residing in brain organoids **(Fig. 4a-c).** In infected organoids (72hpi), this uptake was increased by threefold **(Fig. 4d**), indicating that SARS-CoV-2 infection can perturb the integrity of neuronal circuits by promoting excessive microglial synapse engulfment. Notably, the observed synapse loss (as well as neuronal apoptosis) greatly exceeded neuronal infectivity.

**Figure 4.**
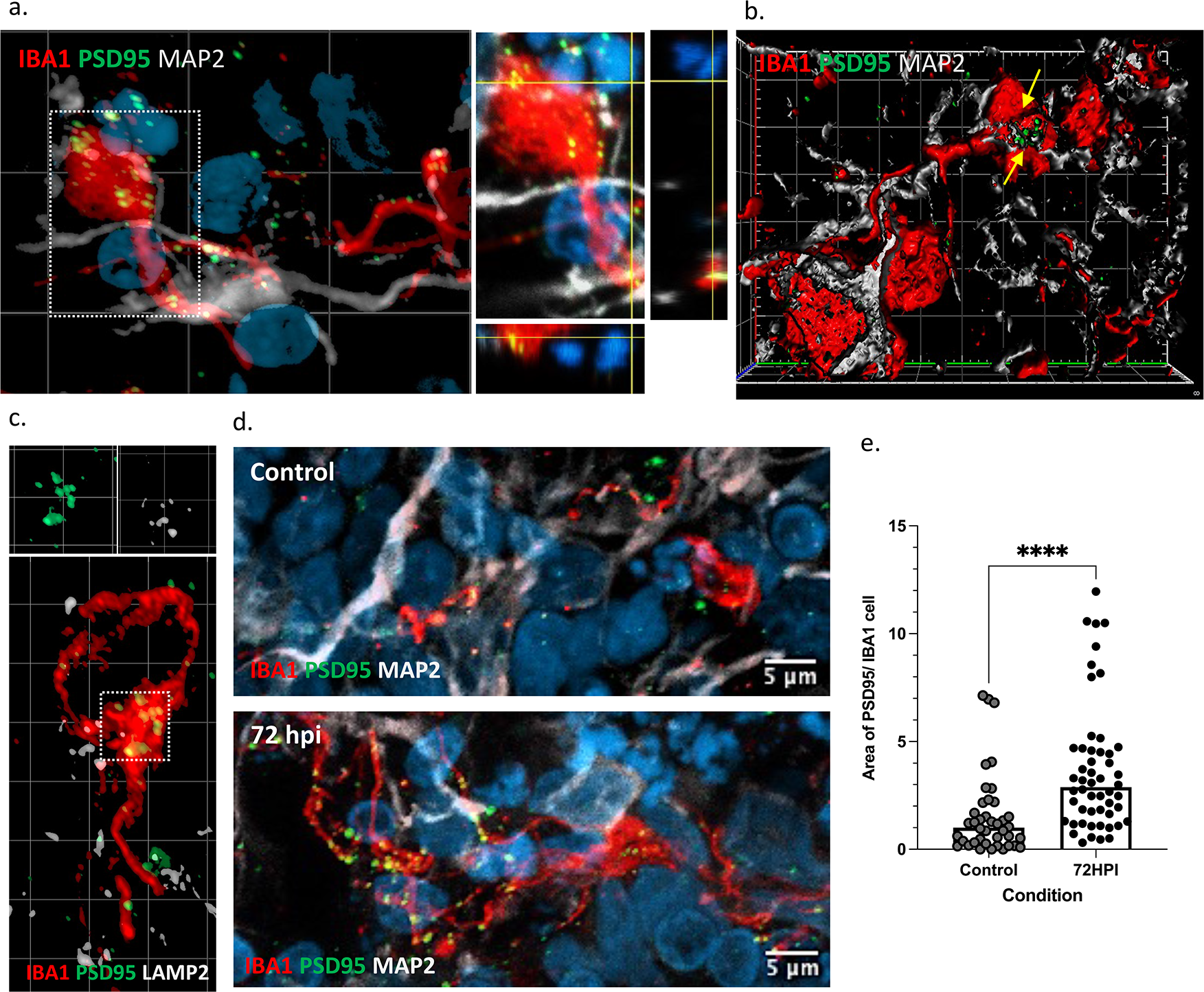
Figure Microglial engulfment of synaptic material in response to SARS-CoV-2 exposure. Super resolution imaging showing (a) microglial processes (IBAl^+^) in proximity to neurons (MAP2\ engulfing post-synaptic material (PSD95^+^) with orthogonal projections showing PSD95 overlapping with IBAl **(b)** 3D reconstruction and surface rendering demonstrating volumes of PSD-95^+^ puncta inside lbal^+^ cells. **(c)** postsynaptic material (PSD95^+^) colocalizing with lysosome associated membrane protein, LAMP2, within phagocyting microglia (IBAl^+^). **(d)** Microglial engulfment of post-synaptic material (PSD-95^+^) in organoids at 72hpi is increased threefold (Mann-Whitney U-test: P<0.0001). Images are representative for each condition. In the graph, bars display medians, and the data is normalized to median uptake in control organoids (n=3 organoids per condition). Each data point represents the mean of all cells per field of view.

### Single cell characterization of brain organoids containing resident immune cells

Droplet-based encapsulation of single cells using the 10X Genomics chromium system allowed us to profile and compare transcriptomes of individual cells isolated from SARS-CoV-2 infected brain organoids as compared to uninfected control organoids. Given the extent of cell death in the organoids exposed to an estimated MOI of 0.3, we then decreased the MOI to 0.1. Since microglial numbers were low in 56 DIV organoids, we also used organoids cultured up to 110 DIV and enriched for CD11b^+^ cells (MACS). In this way we were able to obtain good-quality transcriptomic data from fresh cells isolated in three experimental conditions: uninfected control organoids (7257 cells), 24hpi organoids (15254 cells), and 72hpi organoids (2825 cells), respectively **(Supplementary Fig. 2b).** Then, we integrated pre-processed data from each condition and performed unsupervised graph-based clustering to obtain cellular clusters with similar transcriptomic identities across conditions **(Fig. 5a, Supplementary Fig. 2a-d).** Supervised inspection of top differentially expressed genes (DEGs) per cluster combined with cell type-specific gene signatures and cluster correlations to other developing human brain^25–31^, as well as organoid^32–34^, datasets confirmed 16 clusters **(Fig. 5b-d, Supplementary table 1, Supplementary Fig. 3, 4a-e).** To address if organoid grown microglia at 110 DIV resembled adult or fetal primary microglia, we performed integration of single cell RNA sequencing data from our microglia cluster with two primary fetal microglia dataset^25,29^, as well as a primary adult dataset ^30^, and concluded that the overall transcriptomic profile was closest to fetal microglia **(Supplementary Fig. 4f).**

**Figure 5.**
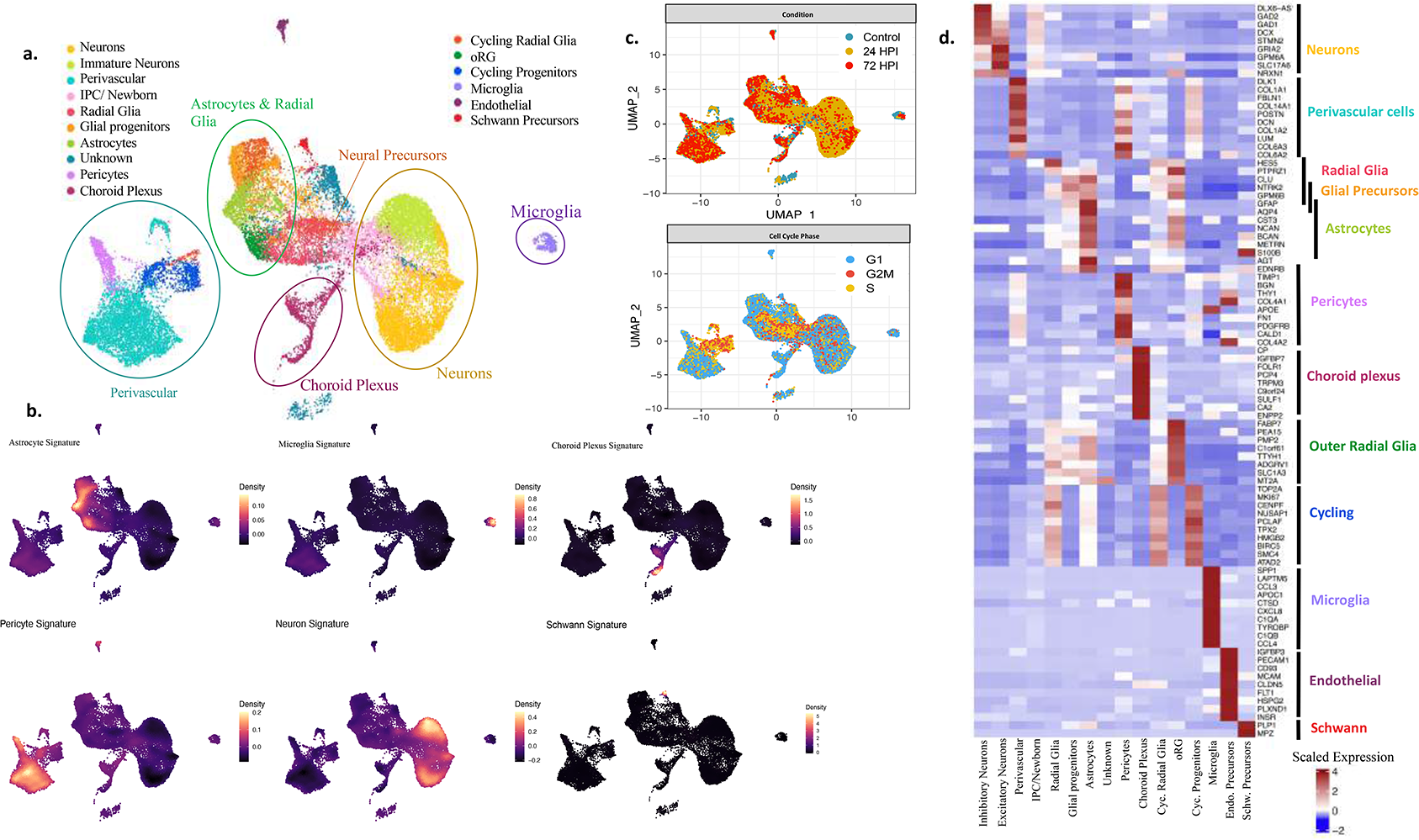
Single cell transcriptome profiling of brain organoids with innately developing microglia. Brain organoids (n=3) per experimental condition (control, 24 and 72hpi) were dissociated into whole-cell suspensions and single cell RNA sequencing libraries were generated using the droplet-based 10X platform. (a) UMAP plot of integrated dataset containing 25336 single cells from all three conditions depicting the presence of key neurodevelopmental cell types. Individual dots representing single cells are colored by the identified cell type. **(b)** Cell type classification obtained by scoring cells by their expression of known individual cell type signatures **(Supplementary table 1)** plotted as estimated joint density on a UMAP.(c) UMAP plot of overall embedding of cells colored by the experimental condition (top) and cell cycle phase (bottom) **(d)** Heatmap showing expression of top-differentially expressed markers (rows) across all clusters (columns) (BH adjusted pvalue<0.05). A list of all differentially expressed genes conserved across conditions for each cluster is provided in supplementary table1.

### Confirming the expression ofSARS-CoV-2 entry factors

We then assessed the expression of previously identified entry factors for SARS-CoV-2 in the generated single cell data and observed that the overall basal RNA expression was relatively low **(Fig. 6a).** However, to some extent, we may have at least partly underestimated *ACE2* expression by enriching for viable cells displaying low tropism and low *ACE2* expression, or not thoroughly capturing *ACE2* expression at the present sequencing depth. In line with previous reports, cells belonging to the choroid plexus showed relatively higher *ACE2* expression^18,35^, and we also observed a more pronounced expression in a subset of neurons with midbrain dopaminergic markers **(Fig. 6a, Supplementary Fig. 4g).** However, this subset of neurons did not exhibit any differential response in relation to other neuronal clusters across conditions **(Supplementary table 3).** Consistent with an underestimation of *ACE2* expression due to e.g., negative selection based on viability, we observed more robust and heterogenous neuronal ACE2 staining using IHC **(Supplementary Fig. 4h).** Similar to *ACE2* expression, proteases involved in viral S protein priming ^39^ *(TMPRSS2* and *TMPRSS4)* also displayed low expression, whereas *FURIN* ^36^ and infection potentiating factor (Neuropilin 1: *NRP1,)* had a wider cell type distribution which included astrocytes, perivascular cells, endothelial cells and microglia **(Fig. 6a).** Microglia also expressed other predicted entry factors *(CTSL* and *CTSB,* Cathepsin L and B, purported to act as substitutes for TMPRSS2 ^37^ **(Fig. 6a).** Next, we identified infected cells by aligning cellular viral transcripts to the whole SARS-CoV-2 genome in the infected conditions. The percentage of infected cells was low (0.1-0.2% of sequenced cells), which was expected given the lower MOI and to some extent removal of non-viable infected cells, although we were able to qualitatively identify infected neurons, radial glia, astrocytes, and choroid plexus cell types **(Supplementary table 2).** Amongst the infected cells, we observed a preferential infection of neurons (p=0.009) and radial glia (p=0.027). For further downstream analyses, we did not make any distinction between infected and non-infected cells, instead focusing on overall cellular responses in the infected conditions.

**Figure 6.**
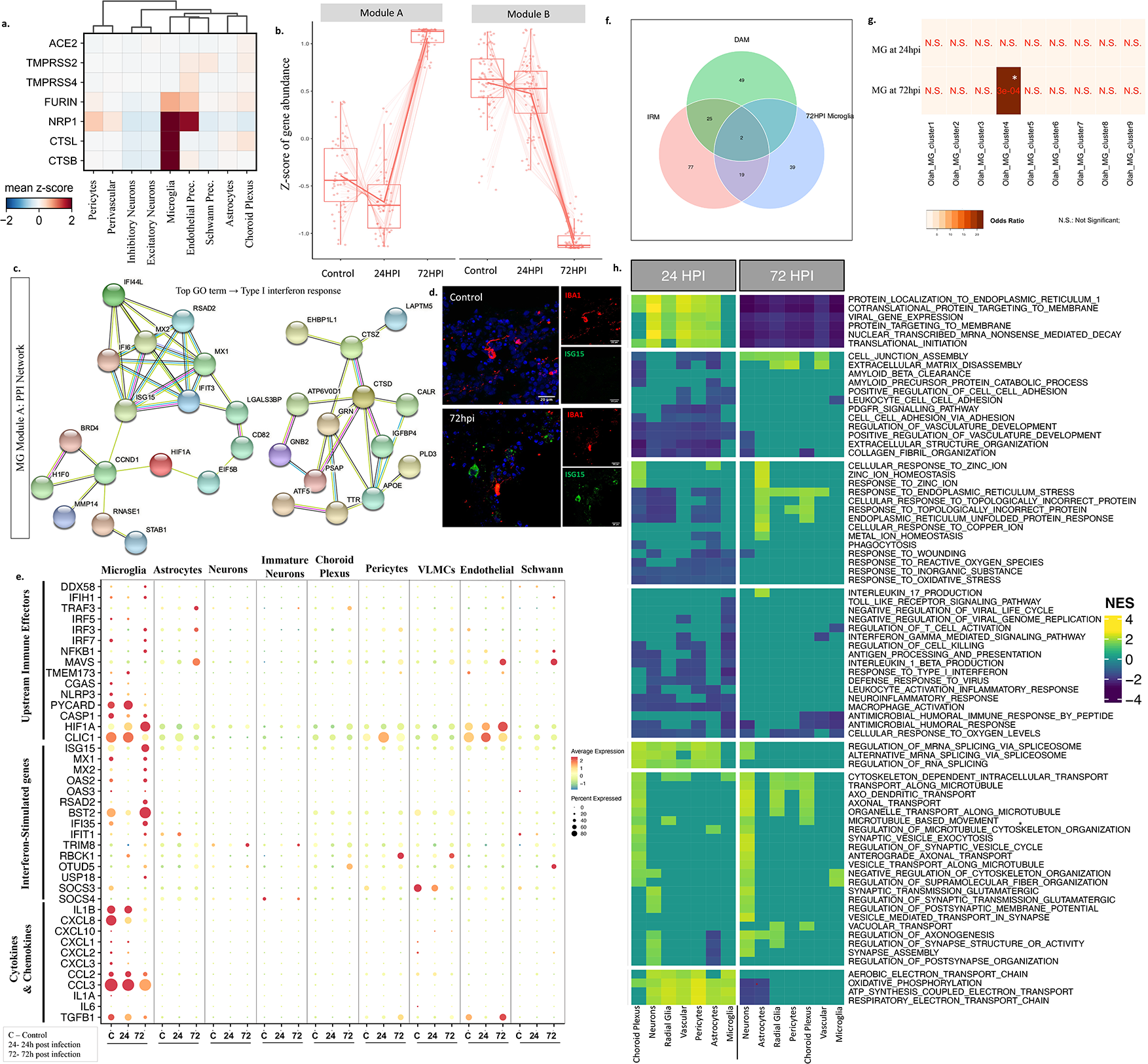
Expression of SARS-CoV-2 entry factors in brain organoids and microglial responses to SARS-CoV-2. **(a)** Heatmap showing relative mRNA expression levels of proposed cellular entry factors utilized by SARS-CoV-2 virus across cell types. **(b)** Modules of genes with similar expression behavior across conditions in microglia identified by pseudo-bulk differential gene expression analysis (using LRT implemented in DESeq2). **(c)** Protein-protein interaction (PPI) network of genes belonging to module A using STRING database with the top Gene Ontology term identified as ‘Type I interferon response’ (Adjusted p-value<0.05).(d) IHC staining validating ISG15 expression in control and 72hpi organoid. Quantifications provided in supplementary fig 5j (e) Dot plot showing expression levels of neuro-immune related genes such as cytokines & chemokines, interferon-stimulated genes, and their upstream effector molecules across infected conditions for each cell type [24-24 hpi; 72-72 hpi]. Color scale represents average log-scaled expression values across all cells, in each cluster. Size of the dot represents the percentage of cells in each cluster, expressing the gene. **(f)** Venn diagram of genes overlapping between gene signatures of Disease-associated microglia (DAM), injury-associated microglia (IRM) and SARS-CoV-2 infected microglia at 72hpi (* BH adjusted p-value < 0.005). (g) Heatmap of comparisons between SARS-CoV-2-associated microglia at 24hpi & 72hpi along with 9 human microglial clusters defined by *Olah et al., 2020,* colored by odds ratio (with p-values shown in red) from hypergeometric gene overlap testing (Fischer’s exact test, Bonferroni correction, adjusted p-value<0.05). Transcriptomic gene signatures were defined as differentially upregulated genes with log2FC > 0.25 and FDR<5%. **(h)** Heatmap showing gene set enrichment analyses *(MSigDB* GO:BP gene sets) of differentially expressed genes (DEGs) from 24hpi versus control (left) and 72hpi versus control (right) for each cell type. Pathways commonly dysregulated across most cell types were selected. All significantly altered pathways for individual cell types are listed in Supplementary table 2 (FDR<5%, Benjamini-Hochberg (BH) correction, NES-Normalized Enrichment Score).

### Interferon responsive microglia with upregulation of genes promoting synapse elimination

To define microglial responses to SARS-CoV-2 exposure, we first performed pseudo-bulk differential gene expression testing across the three experimental conditions, i.e., uninfected control organoids, and organoids 24hpi and 72hpi. An unbiased hierarchical clustering approach generated two unique groups of DEGs (labeled module A and B). Module A comprised of genes whose expression on average decreased at 24hpi and then at 72hpi displayed a more pronounced increase, while module B genes on average displayed reduced expression at 24hpi and a more dramatic decrease at 72hpi **(Fig. 6b).** While module B was largely comprised of genes encoding for cytosolic ribosomal proteins **(Supplementary Fig. 5e**), we detected several interferon-stimulated genes (ISGs) in module A such as *ISG15, MX1, MX2, RSAD2* and *BST2* **(Fig. 6c, Supplementary table 4).** Accordingly, IHC validation of ISG15 protein expression showed a general increase in interferon response in organoids at 72hpi as well as a specific upregulation in IBA1^+^ cells, compared to control organoids **(Fig. 6d, Supplementary Fig. 5j).** Nonetheless, we detected no clear upregulation of pro-inflammatory cytokines in microglia, at either 24hpi or at 72hpi, although several pathogen sensors and upstream effectors of interferon signaling were upregulated including nuclear factor kappa B *(NFKB1)* **(Fig. 6e).** At 24hpi, we detected upregulation of pathways (such as oxidative phosphorylation, release of calcium into cytosol, monovalent inorganic ion transport, lysosomal disruption and hypoxia; **Supplementary Table 3)** that can serve as activating stimuli for primed inflammasome components *(NLRP3, PYCARD, CASP1),* and are required for the production and maturation of pro-inflammatory cytokines (**Fig. 6e**)^38^. Taken together, this suggests that SARS-CoV-2 activates interferon signaling in microglia although the expression of pro-inflammatory cytokines are at normal levels already at 24hpi.

In the infected conditions, we in unbiased analyses observed a significant upregulation of pathways related to neurodegenerative diseases, including Alzheimer’s disease (AD) and Parkinson disease (PD) **(Supplementary Fig. 5f).** Notably, both these disorders are characterized by a microglia-mediated early synapse loss^39^, and display an increased incident risk after Covid-19 infection^3^. This led us to examine the microglial transcriptional state more closely after exposure to SARS-CoV-2 in relation to previously described disease-associated microglial activation states. First, we compared SARS-CoV-2-exposed microglia to such activation states observed in experimental murine models. Injury responsive microglia (IRM) have been described in mice close to experimentally induced focal demyelination (7 days post lesion) and are to a large extent defined by upregulation of ISGs ^40,41^, while the disease-associated microglia (DAM) described in mouse models of AD and ALS are thought to restrict neurodegeneration and disease development^42^. SARS-CoV-2 exposed microglia then displayed a significant enrichment for genes associated with the IRM activation state (p=6. 9×10^−24^, **Fig. 6f**), and less so for the unique DAM signature genes (p=0.11), while still displaying upregulation of genes implicated in AD such as *APOE, CTSZ* and *GRN* with a non-significant decrease of core microglial homeostasis genes *(P2RY12, CX3CR1, P2RY13, CD33, TXNIP;* **Supplementary Fig. 5g-h).** In fact, *APOE* and *CTSZ* were observed to be commonly dysregulated across all three signatures (IRM, DAM and SARS-CoV-2 exposed microglia), suggesting unique as well as shared activation mechanisms across these microglial states. Further, we compared the SARS-CoV-2 microglial transcriptomic signature to previously identified human microglial populations identified in autopsy and surgical brain tissues obtained from subjects in the Memory and Aging Project (MAP) ^43^ and found an enrichment with genes in so-called interferon responsive microglia (cluster 4 in this dataset) **(Fig. 6g**), defined by *ISG15* and with increased expression of multiple sclerosis (MS) and AD susceptibility genes. To exclude effects of tissue processing, we also compared microglia from the organoids with a cluster defined by general cellular distress due to tissue processing (cluster 3) but observed no enrichment **(Fig. 6g).**

Actin-cytoskeletal remodeling pathways, essential for promoting migration and phagocytosis, ^44^ were also upregulated in microglia exposed to SARS-CoV-2 **(Fig. 6h, Supplementary Fig. 5f).** Consistent with this upregulation in microglia, and the extensive neuronal cell death and decreased postsynaptic density in infected organoids, neurons at 72hpi downregulated the expression of ‘don’t-eat-me’ signals such as *CD46* and *CD200* **(Supplementary Fig. 5i**)^45^. Consequently, we observed increased microglial expression of genes associated with microglia-mediated phagocytosis ^46–49^, such as *CD68, TREM2, ITGB5, CD47, MSR1, CALR,* as well as genes previously identified to be directly involved in synapse elimination in the aftermath of viral encephalitis, such as *C3, C3AR1* and *FCGR3A*^8^ **(Supplementary Fig. 5h).**

In summary, this suggests that microglia in response to SARS-CoV-2 exposure quickly adopt transcriptional states that are unique but also overlap with signals observed in neurodegenerative diseases, and that these states are likely to include both protective - e.g., clearing of debris and defense, as well as detrimental - e.g., inducing excessive neuronal apoptosis, and synaptic stripping responses.

### Astrocytic subclusters with enrichment for genes implicated in neurodegenerative diseases

In response to SARS-CoV-2, astrocytes displayed DEGs enriched for mechanisms involved in cell cycle, carbon metabolism and disorders such as Huntington’s disease (HD) and PD **(Fig. 7a).** Already at 24hpi, three subclusters of astrocytes could be identified: AS-0, AS-3 and AS-4 **(Fig. 7b).** AS-0 cells expressed relatively higher levels of anti-viral ISGs such as *IFIT1, ISG15,* and *SCRG1,* while AS-3 and AS-4 cells had a proliferative profile *(TOP2A,* MK167) and exhibited lower expression levels of *GFAP* **(Fig. 7b-d).** Subcluster AS-2, comprising of cells sampled at 72hpi, showed elevated levels of *GFAP* and *STAT3,* indicative of reactive astrogliosis^50^, along with phagocytosis related genes such as *MEGF8* and *ABCA1* **(Fig. 7c).** We also observed upregulation of metal ion (zinc, copper, iron) homeostasis pathways involving metallotheionins *(MT2A, MT1X, MT1E)* in AS-2 astrocytes at 72hpi **(Fig. 7d).**

**Figure 7.**
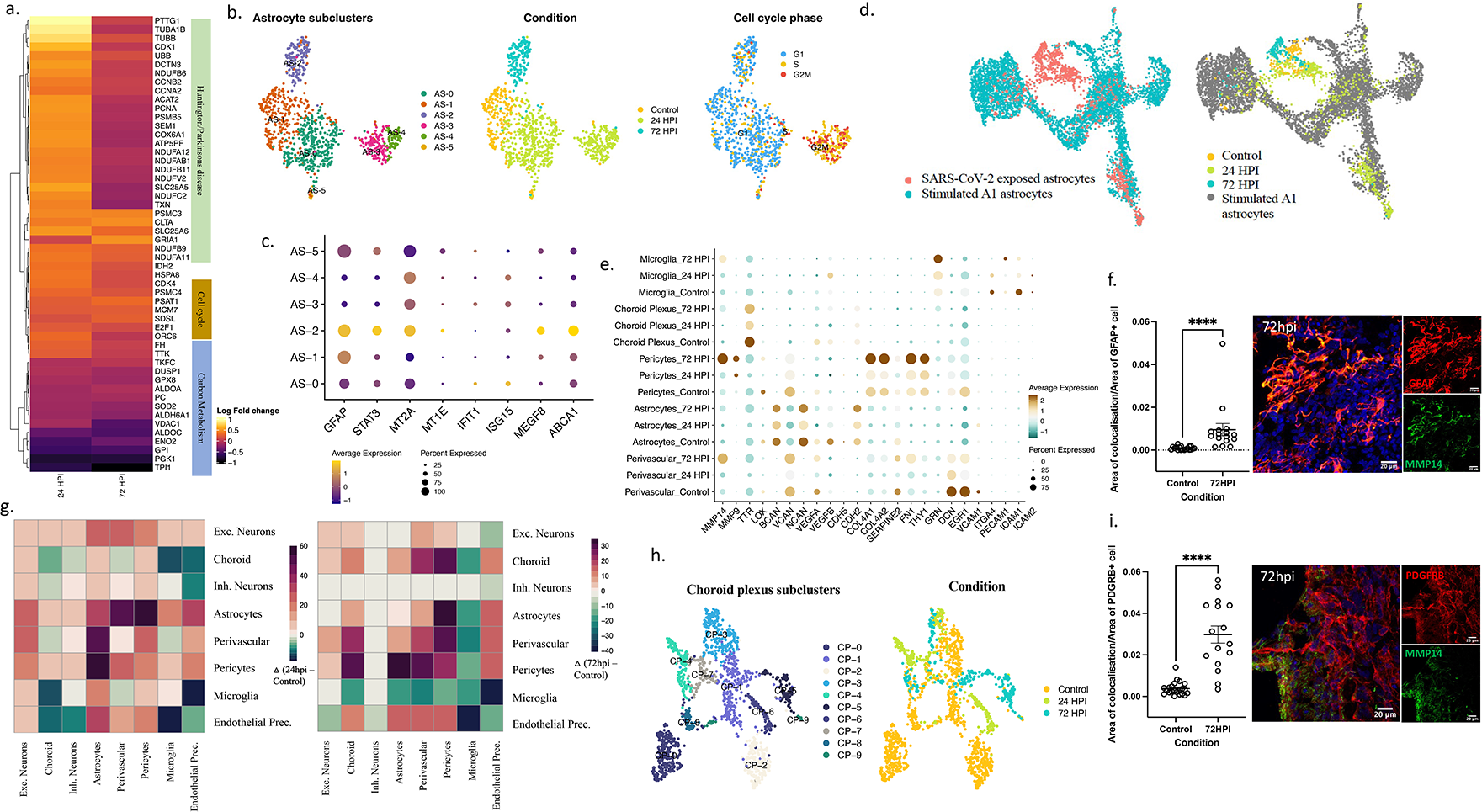
Temporal heterogeneity in astrocytes and choroid plexus responses to SARS-CoV-2. **(a)** Differential expression analysis of SARS-CoV-2 exposed astrocytes versus control astrocytes, highlighted DEGs in a heatmap with top implicated pathways using overrepresentation test (FDR<0.05, Benjamini-Hochberg correction; Scale bar-Log2fold change values). **(b)** Cells belonging to the astrocyte cluster, as shown in Fig 4A, were extracted and subjected to further unsupervised clustering. UMAP plot of astrocytes shows 6 unique subclusters (AS-0, AS-2, AS-2, AS-3, AS-4 & AS-5; left) with cells colored by condition (middle) and cell cycle phase (right) (clustering was performed upon regressing cell cycle effects). Each dot represents a single cell colored by the respective groups shown on the right side of each plot. (c) Dot plot of scaled average expression of genes involved in astrocytic engulfment across astrocyte subclusters (adjusted P<0.05) **(d)** UMAP plot of single-cell transcriptomic data of human iPSC-derived Al-astrocytes from *Barbar et al., 2020,* integrated with astrocytes from our study using canonical-correlation based-approach (CCA) (implemented in Seurat). Cells are colored by their respective dataset and experimental condition. **(e)** Average-scaled expression of genes related to blood-brain-barrier integrity and function for different cell type clusters across control and infected conditions. (f) IHC staining and corresponding quantification validating increased MMP14 expression in GFAP^+^ cells and (i) PDGFRB^+^ cells. **(g)** Mapping intercellular interactions across cell types upon infection. Heatmap showing changes in the number of interactions between cell types at 24hpi (left) and 72hpi (right) resulting from SARS-CoV-2 infection. Ligand-receptor interactions were inferred in the single-cell transcriptomic data using *CellphoneDB.* Green depicts a negative difference indicating loss of interactions whereas purple depicts a positive change indicating gain of interactions between the cell type pair when compared to the control. (h) UMAP plot of all Choroid plexus cells colored by their identified subclusters (left) and experimental condition (right).

In neurodegenerative diseases, astrocytes have been shown to express markers of a putative reactive neurotoxic state commonly referred to as A1^51^. Given that astrocytic DEGs from infected organoids were enriched for such diseases, we integrated single-cell transcriptomic data from Al astrocytes (stimulated with IL-lA, TNF-α and Clq)^52^ and SARS-CoV-2 exposed astrocytes via canonical correlation analysis (CCA). Although the majority of SARS-CoV-2 exposed astrocytes (AS-0, AS-1, AS-2) clustered separately from Al astrocytes, the proliferative clusters (AS-3 and AS-4), as well as a subset of AS-0 astrocytes, showed clustering similarities with a portion of Al astrocytes **(Fig. 7c).** Furthermore, higher expression levels of metallotheionins, as in AS-2 cluster (72hpi), have been observed in astrocytes from HD patients^53^. In summary, similar to the observed effects in microglia, we observe responses in astrocytes to SARS-CoV-2 exposure that share features with detrimental cellular states observed in neurodegenerative diseases, but also a strong temporal heterogeneity that includes unique cellular states.

### Secretome alterations indicating a compromised capacity to keep BBB integrity

Potential entry routes to the CNS for SARS-CoV-2 includes the blood-brain barrier (BBB) or the blood-cerebrospinal-fluid-barrier (BCSFB), involve cell types such as endothelial cells, pericytes, astrocytes and choroid plexus-epithelium. Upon encountering pathogens, these barrier cells act in concert to activate and regulate signal transduction pathways that aid invasion of peripheral immune cells and restoring of CNS homeostasis^54^. Correspondingly, at 24hpi we found increased ligand-receptor communication between astrocytes, neurons, microglia, perivascular, and endothelial cells, whereas at 72hpi, both microglia and neurons significantly reduced their communication with most other cell types, while choroid plexus cell types gained interactions **(Fig.** 7g). Sub-clustering of choroid plexus related cells **(Fig.** 7h) showed two control enriched clusters (CP-0 and CP-2), consisting of genes involved in maintaining solute homeostasis *(AQP1),* barrier or extracellular matrix (ECM) integrity *(TJMP1, LGALS1, COL1A2, COL2A1, COL3A1)* and amyloid clearance (7TH, *MT3),* amongst others (Supplementary **Fig.** 7c). At 24hpi, we observed an enrichment of metallotheionins *(MT1M, MT1E, MT1X, MT1F, MT2A:* CP-6), similar to what we observed in astrocytes at 72hpi, whereas genes involved in viral defense response *(SAMHD1),* regulation of reactive oxygen species (ROS) *(50D3)* and ECM organization *(FBLN1, SYNE1)* were enriched at 72hpi (CP-5) **(Supplementary Fig. 7c).** In addition to choroid plexus, differential expression analysis showed significant increase in expression of matrix metalloproteases and ECM-regulatory enzymes *(MMP14, MMP9, LOX)* at 72hpi in microglia, astrocytes and perivascular cells, whereas endothelial cells reduced expression levels of solute carrier *SLC2A1* **(Fig. 7e**), all in agreement with signaling that promotes compromised CNS barriers. Accordingly, by IHC we further validated increased MMP14 protein expression by PDGFRB^+^ (pericytes) as well as GFAP^+^ cells within 72hpi organoids **(Fig 7f-i).** Further, astrocytes, pericytes and choroid plexus related cells were observed to downregulate VEGF signaling, a potent inducer of BBB permeability, at 24hpi but then at 72hpi instead display an upregulation, coinciding with increased signaling for leukocyte chemotaxis and activation in astrocytes, as well as antigen processing and presentation in choroid plexus **(Fig. 7e).** In aggregate, this suggest that cell types implicated in barrier functions change their communication upon SARS-CoV-2 infection, indicative of compromised integrity, as well as altered secretome and signaling with peripheral immune cells.

### Metabolic dysregulation across cell types

Protein accumulation in the endoplasmic reticulum (ER) following viral infection or excessive production of secretory proteins can induce ER stress and initiate counter measures in the form of reduced translation and export to ER along with unfolded protein responses (UPRs). ^55^ In exposed organoids, we detected an upregulation of pathways related to UPRs (IRE1, PERK), ER stress, proteasomal degradation and autophagy, among cell types such as microglia, astrocytes, choroid plexus and perivascular cells **(Supplementary Table 3).** Further, a partial translational inhibition could be observed in microglia (module B, as discussed before), as well as most of the other cell types **(Supplementary Table 3, Fig. 5h).** We also observed a downregulation of the ROS metabolism across cell types, which can lead to dysregulation of the antioxidant cellular systems and result in oxidative stress^56^. SARS-CoV-2 then relies on excessive glucose for its replication, ^57^ and shunts glucose from glycolytic pathways to fuel oxidative phosphorylation and ATP production in a process known as the Warburg effect^58^. In our dataset, pathways pertaining to cellular respiration (oxidative phosphorylation, electron transport chain coupled to ATP synthesis) were enriched at 24hpi in neurons, while those of glucose and pyruvate metabolism were downregulated **(Supplementary Table 3).** Furthermore, transcription of glucose transporter, GLUT1 *(SLC2A1),* lactate dehydrogenase *(LDHA)* along with monocarboxylate transporter, MCT4 *(SLC16A3),* was downregulated in astrocytes at 24 and 72hpi, indicating deficits in glucose export and redox cycling necessary for neuronal energization^59^ **(Supplementary Table 3).** This suggests that following exposure to SARS-CoV-2, CNS cell types undergo major stress that results in metabolic changes which are likely to impact cellular communication such as the important interplay between astrocytes and neurons.

## DISCUSSION

Long-lasting neurobehavioral and cognitive impairments have been increasingly recognized in the ongoing Covid-19 pandemic^6,7,60^. In the following 6 months after recovery, even non-hospitalized patients display an increased risk of receiving a neurological or psychiatric diagnoses^6^ while imaging studies have revealed widespread micro-structural disruptions in a subset of the recovered patients^5,6^ To a large extent, the described sequalae overlap with post-infection syndromes described in patients that has recovered from infection with more uncommon neuro-invasive RNA viruses, such as the West Nile virus or the Zika virus ^8^ In murine models for encephalitis caused by these viruses, CNS-damage continues after virus elimination ^8^ and can be largely attributed to interferon-responsive microglia that excessively eliminate pre- or post-synaptic termini as well as induce neuronal apoptosis ^9,10^. Here we infect brain organoids containing innately developing microglia with live SARS-CoV-2 virus and observe a widespread neuronal cell death as well as a clear decrease in post-synaptic density. We show that microglia in the infected organoids increase the engulfment of postsynaptic termini with a concomitant upregulation of interferon-responsive genes as well as genes promoting phagocytosis and synapse elimination, while neurons down-regulate ‘don’t-eat-me’ signals. Transcriptomic profiling on single cell level reveal that microglia in the infected condition largely overlap with the signature of interferon-responsive microglia described in AD or PD. Interestingly, these disorders are also characterized by an early synapse loss driven by microglial synapse elimination^39^ and the incidence is increased post COVID-19^3^. Similar, microglia-mediated synapse elimination has also been observed in schizophrenia models ^61^, another disorder that display an increased incident risk after COVID-19 infection^3^.

SARS-CoV-2 infection in brain organoids also leads to secretome alterations in astrocytes, pericytes and choroid plexus related cells indicating a compromised capacity to keep BBB integrity, as well as an upregulation of pathways related to signaling with peripheral immune cells. The murine models of Zika or West Nile encephalitis have revealed a role of infil trat ing CD8^+^ T-cells to activate interferon signaling **in** microglai ^10^, then suggesting that a similar mechanism could exacerbate synaptic stripping secondary to SARS-CoV-2 exposure **in** a model that includes CD8^+^ T-cells.

Although it is unclear to what extent the used virus loads recapitulate *in vivo* exposure, it is noteworthy that we here observe the microglial responses on modest virus titers that cause a limited infection of neuronal cells. While viral gene copies increase in the supernatants of infected organoids, the number of PFUs promptly decrease at 72hpi. This suggests that the limited number of infected cells either have an inefficient assembly and shedding of viral particles, or that cell death decreases the availability of viable cells to sustain the production of fully assembled viral particles.

Existing protocols for generating brain organoids have several constraints that also limit our interpretations. Importantly, brain organoids most closely resemble the developing fetal brain rather than the mature adult brain. Thus, we cannot exclude important differences exist in the cellular responses and tropism between immature and more mature brain cells, and primarily our model should be considered to recapitulate the responses to SARS-CoV-2 of the developing brain. Finally, our model does not include peripheral immune cells and thus cannot capture effects related to infiltrating immune cells. For example, a likely scenario is that infiltration of CD8^+^ T-cells would have amplified microglial responses. However, by excluding infiltrating monocyte-derived cells, the model is optimized for assessing microglial responses thus avoiding difficulties with distinguishing them from that of peripheral macrophages.

In summary, we here provide an experimental approach for modeling viral encephalitis that includes tightly orchestrated responses of microglia and astrocytes in the context of neuronal circuits. Challenging this model with modest live SARS-CoV-2 virus titers, we observe extensive neuronal cell death and microglia-mediated synapse loss. A key next step will be to determine the clinical importance of such molecular processes, and specifically whether they may contribute to synapse loss and the neurocognitive and neuropsychiatric symptoms observed in a subset of COVID-19 patients across different developmental stages. If so, these microglia-containing organoid models may provide an opportunity to evaluate microglia-targeted therapeutics aimed at minimizing or preventing these COVID-19 sequelae.

## MATERIALS AND METHODS

### Ethics

All individuals signed a written informed consent before participating in the study, as approved by the Institutional Review Board of Partners HealthCare (Boston, MA, USA) and the Regional Ethical Review Boards in Stockholm, Sweden. All relevant ethical regulations were followed when performing the study.

### Vero E6 cells

Vero E6 (ATCC-CRL-1586) cells were maintained in Dulbecco’s modified eagle medium (DMEM, Cytiva) supplemented with 5 % heat-inactivated fetal bovine serum (FBS, Cytiva), 100 U/mL penicillin and 100 μg/mL streptomycin (Cytiva).

### iPSC reprogramming

Two healthy human iPSC lines (males) were obtained from the MGH Neurobank. Briefly, iPSC colonies were obtained using mRNA reprogramming in a feeder-free culture system as described previously ^24^. Stable iPSCs were expanded in mTESR plus media (STEMCELL Technologies) and on biolaminin 521 LN-coated (BioLamina) plates. iPSCs were then purified using MACS with anti-TRA-1-60 MicroBeads (Miltenyi Biotec) on LS columns. All fibroblasts and iPSCs were screened and found negative for Mycoplasma and stained positive for pluripotency markers like octamer-binding transcription factor 4 (POU domain, class 5, transcription factor 1) and TRA-1-60.

### Brain organoid cultures

Undirected brain organoids were prepared from single cell suspension of human iPSCs as previously described^21^. Briefly, embryoid bodies (EBs) were generated by seeding 9000 single cells in each well of a low attachment 96-well U-bottom plate with Y-27632 ROCK inhibitor (10 μM) for one day. EB media was replaced on day 5 with Neural induction media in the same 96-well plate. On day 10-11, EBs were embedded in 30μl Matrigel (Corning) using sheets of dimpled parafilm and incubated for 20 min at 37°C as previously detailed. Matrigel embedded single EBs were then either transferred to a 24-well plate in 3ml of expansion media per well, on an orbital shaker (90 rpm), or to a 12-well miniaturized bioreactor (largely overlapping with the design proposed by Qian et al.)^62^, until further use.

### Induced microglia-like cells

iMGs were derived from monocytes (donated from one healthy male) using established methods previously described in detail ^23,24^. Briefly, whole blood was collected into vacutainer cell preparation tubes containing sodium citrate as an anticoagulant (Becton, Dickinson and Company) and processed as per the manufacturer’s instructions. Peripheral blood mononuclear cells (PBMC) were isolated, washed twice with PBS by centrifugation and suspended in heat-inactivated fetal bovine serum (FBS; Sigma) containing 10% DMSO (Sigma). The cell suspension was then divided into aliquots and cryopreserved. Generation of induced microglia-like cells (iMG) from PBMCs were carried out using methods previously descri bed^23^ with minor modifications. Briefly, cryopreserved PBMC samples were transferred from liquid nitrogen freezer to a 37 °C water bath. Once the cell suspension had been thawed, it was gently pipetted into 10 ml of pre warmed complete RPMI medium (CM) consisting of basal RPMl-1640 supplemented with 10% heat-inactivated FBS and 1% penicillin-streptomycin (P/S; Thermo Fisher Scientific). Following centrifugation (300 *g* for 5 min at room temperature), the supernatant was discarded, and the cell pellet was resuspended in the appropriate volume of CM. Isolated PBMCs were cultured at a density of 5×10^5^ cells/1 ml CM on 24-well plates coated with Geltrex (Thermo Fisher Scientific). After 24 h of incubation, the media was replaced with RPMl-1640 supplemented with lx glutamax (Life Technologies), 1% P/S and 0.l l⍰lμ gl⍰lml-^1^ of interleukin (IL)-34 (R&D Systems) and 0.01 μg ml ^−1^ of granulocyte macrophage colony-stimulating factor (GM-CSF; R&D Systems). After 7 days, fresh media was added. At day 11, cells were used for infection.

### Virus isolate

SARS-CoV-2 (GenBank: MT093571.1), originally from the Public Health Agency of Sweden, was propagated on 90% confluent Vero E6 cells for 3 days at 37 °C. Cell debris were removed by centrifugation at 300 RCF for 5 minutes and the viral supernatant was aliquoted into cryovials and stored at −80 °C. Viral titers were quantified by PFU assay.

### Viral infection

All experimental studies involving infectious SARS-CoV-2 were performed within the biosafety level 3 (BSL3) facility at Karolinska Institute. From single cell dissociation experiments, approximately l×10^6^ cells were found to be present in a D56 organoid. Based on this, 3×10^5^

PFU/ml of SARS-CoV2 was used to infect organoids for 2h (such that the estimated MOI was 1 for periphery and 0.3 for the whole organoid) in a spinning bioreactor. Following viral exposure, the organoids were washed twice with PBS and transferred into fresh medium (2ml) to follow the course of infection. To monitor viral replication, 100μ1 of medium was collected at specified time points, centrifuged at 300g for 5 min and the supernatant was collected in trizol for qPCR analysis. For single cell RNA sequencing (scRNA-seq), the viral load was reduced to 1 ×10^5^ PFU/ml (MOI 0.1) in order to reduce the percentage of cell death observed with the previous MOI

### Plaque forming unit assay

PFU assays was performed on 24-well cell culture plates seeded with 2×10^5^ Vero E6 cells per well. A serial dilution of the inoculum medium (consisting of either the viral stock or experimental samples in DMEM) was prepared and 200μl used to infect each well. The plates were then incubated for 1 hour at 37 °C, 5% CO2. After the incubation period, the wells were washed twice with PBS and 1 ml of overlay medium (2:3 mix of 3% high density carboxymethyl cellulose and complete DMEM medium) was added to each well and the plates incubated for 3 days at 37 °C, 5% CO2. The plates were subsequently inactivated with 1 ml of 10% formaldehyde solution overnight at RT, washed twice with PBS, stained with 200μl of crystal violet solution for 30 min at room temperature and the plaques counted.

### lmmunohistochemistry

Organoids were fixed with 4% paraformaldehyde for 20 min at RT followed by 24h in 30% sucrose in PBS. Organoids were then embedded in OCT (VWR) and frozen at −80°C. Cryosections (16 um) of brain organoids were incubated with blocking solution for lh (10% of Normal Donkey serum in 0.3% Triton X) in 1xPBS. Blocked cryosections were incubated with the respective primary antibodies, diluted in blocking solution and incubated overnight at 4°C:, anti-b-111-tubulin (mouse, Promega G712A, 1:500), Cleaved caspase-3 (rabbit, Cell Signaling, 1:300), anti-PAX6 (mouse, Developmental Studies Hybridoma Bank, 1:100), anti-SOXl0 antibody (goat, R&D AF2864, 1:50), anti-OLIG2 (goat, R&D AF2418, 1:100), anti-GFAP (mouse, Sigma G3893, 1:100), anti-lBAl (rabbit, Wako 019-19741, 1:100), anti-MMP14 (mouse, R&D MAB918-SP, 1:100), anti-lSG15 (mouse, Santa Cruz Ltd sc166755, 1:100), anti-PSD95 (mouse, Abeam 13552, 1:100), anti-PDGFRB (goat, R&D AF385-SP, 1:200), anti-LAMP2 (goat, R&D AF6228-SP, 1:200) and SARS-CoV-2 (2019-nCoV) Nucleoprotein / NP Antibody (rabbit, Nordicbiosite 158-40143, 1:200). Cryosections were washed three times with lxPBS and incubated with the secondary antibody for lh at RT. Secondary antibodies **(all** conjugated to Alexa Fluor 488, 555 and 647) were purchased from Life Technologies and used at a 1:500 dilution. Then, samples were washed three times with (1xPBS), incubated for 5 minutes with fluorescent nuclear DAPI stain (VWR; 1:5000), and mounted with DAKO immunofluorescence mounting media (Life Technologies). Image acquisition was performed using a Zeiss LSM800 confocal microscope (40X/1.2 W objective). Super resolution imaging acquisition was performed using a Zeiss LSM900-Airy2 scan confocal microscope (63X/1.4 oil objective) for representative images.

### Image quantification

Immunofluorescence images were acquired using Zeiss LSM-800 confocal, LSM900-Airy2 (super-resolution) microscope and analyzed using CellProfiler software to measure and classify cells according to the expression of the selected markers. First, the software was trained to automatically segment the images into: (1) cells (DAPI^+^ objects); (2) NP or casp3 objects, followed by relating objects (1) and (2). For cell type specific quantifications, (2) was related to (3) objects segmented by cell type specific markers stained by antibodies indicated above. Finally, the related objects were expressed as a percentage of DAPI^+^ cells. For validation of DE genes, 40X confocal images were obtained as tiles for the whole organoid and were processed in lmageJ/FIJI software (v2.1) for colocalization analyses.

For quantification of microglial engulfment of post-synaptic termini, images were acquired throughout the organoid while focusing on areas with IBA^+^l cells (blinded to condition). The images were then processed using lmageJ/FIJI software (v2.1) macro consisting of the following steps: duplication of the red channel (corresponding to IBA1); background subtraction (rollingball set to 20); conversion to binary image; filters set to maximum and minimum of radius=5 followed by particles regarded as ROls with size >1000 pixels. Then, for each ROI in a FOV, the green channel (corresponding to PSD-95) was used to set threshold (50, 255) and particles with size >3 pixels (circularity 0.01-1.00) were analyzed for number and area of each synaptic puncta. Total synaptic area in a ROI was normalized by the area of the corresponding ROI to get a measure of engulfment. The same set of images were used to measure area of colocalization between channels corresponding to MAP2 and PSD-95, respectively, followed by normalization with total MAP2 area in a FOV.

### Quantitative PCR (qPCR)

RNA extraction was performed using DirectZol RNA-Miniprep Kit (Zymo Research Inc.), following the manufacturer’s protocol. Samples collected in TRI reagent were applied to Zymo-spin columns. DNase I treatment was performed for samples with genomic DNA. Bound RNA was washed, diluted in 30ul nuclease-free water and stored at −80°C. Quality and concentration of extracted RNA was determined using a NanoDrop (Thermofischer Scientific). Total RNA was reverse-transcribed to cDNA using the High-Capacity RNA-to-cDNA Synthesis kit (Thermofischer Scientific) following manufacturer’s protocol. Reverse transcription reactions were carried out in 20ul reaction volume containing 1ug of input RNA with the following thermal cycler conditions: 37°C (60 minutes) ^+^ 95°C (5 minutes) ^+^ 4°C (∞) . cDNA was further diluted to 1:3 and used as templates for PCR reactions.

PCR reactions were performed using the StepOnePlus™ Real-time PCR system (Applied Biosystems, Thermofischer Scientific) with PowerTrack™ SYBR Green Master Mix. Expression of viral mRNA was assessed by evaluating threshold cycle (Ct) values. Relative expression levels of N gene in cell lysates were normalized against a housekeeping gene, GAPDH (glyceraldehyde-3-phosphate dehydrogenase), according to the Delta delta Ct method. Absolute quantification of viral gene expression levels in the supernatant were performed using the standard curve method. Primer sequences used in this study are:

GAPDH Forward - GGTGGTCTCCTCTGACTTCAACA
GAPDH Reverse - GTGGTCGTTGAGGGCAATG
N gene Forward - CATTGGCATGGAAGTCACAC
N gene Reverse - TCTGCGGTAAGGCTTGAGTT
RdRp Forward - CGCATACAGTCTTRCAGGCT
RdRp Reverse - GTGTGATGTTGAWATGACATGGTC

### Dissociations for single cell RNA sequencing

Briefly, three organoids per condition (control, 24hpi, 72hpi) were washed twice with DPBS without ions and cut into small pieces using a sterile scalpel. Single cell suspension was generated using neural dissociation kit (Miltenyi Biotec) according to manufacturer’s protocol. Microglia was enriched using CD11b magnetic microbeads (Miltentyi Biotec) using MACS. Viability and cell numbers were assessed for both MACS-enriched and flow-through fractions before proceeding with the 10X protocol.

### Single-cell RNA-sequencing

Two single cell suspensions per condition were loaded onto a single Chromium controller chip v3.1 (l0X Genomics), with a target output of 6000-7000 cells per channel. For the six loaded channels, GEM generation, barcoding, cDNA amplification and library preparation was performed using the Single-cell 3′ Gel Bead and Library v3.1 kit (10X Genomics) at the Eukaryotic Single Cell Genomics facility (Scilifelab, Sweden). Amplified cDNA and final libraries were evaluated on a Bioanalyzer for quality control and sequenced together on lllumina NovaSeq 6000 platform.

### Data processing

Sequenced data was processed through the CellRanger Software (5.0, 10X Genomics) and transcripts were aligned to a combined reference of Human GRCh38-3.0.0 and SARS-Cov-2 genome (GenBank: MT093571.1). Filtered feature-barcode matrices from CellRanger were further subjected to quality control measures where low-quality cells with < 200 uniquely expressed genes and high percentage of mitochondrial reads (>20%) were filtered out. Doublets were identified using scDblFinder package^63^ and removed with caution. Genes expressed in fewer than 5 cells were filtered out. Filtered count matrices were merged and analyzed downstream using the Seurat package^64^. Expression data per experimental condition was normalized using the regularized negative binomial regression method implemented in SCTransform^65^, while regressing out the difference between the G2M and S phase scores, which served as a confounding factor in our dataset. Anchors across conditions were identified using FindlntegrationAnchors function and samples were integrated using canonical correlation analysis (CCA) implemented in the IntegrateData function of Seurat. Linear data compression using principal component analysis (PCA) was performed on 3000 highly variable genes. Top 30 PCs were used as the input to perform non-linear dimensionality reduction using UMAP.

### Clustering and cell-type identification

We constructed a k-nearest neighbor (KNN) graph based on Euclidean distance in 30 PCs and performed unsupervised graph-based clustering using the Louvain algorithm (modularity resolution = 0.8) with functions provided in Seurat. Preliminary clustering of 39,808 cells identified 21 clusters yielding a clear separation of neuro-ectodermal clusters. Non-neuroectodermal clusters that did not express known marker genes related to cell types present in the brain, were removed. The remaining 26,148 cells were then subjected to second-level clustering in an integrated space (resolution= 0.6), yielding 16 final clusters. The clustering was visualized using UMAP embedding in two dimensions. Top differentially expressed genes conserved across conditions for the final clusters were identified using a non-parametric Wilcoxon rank-sum test in FindConservedMarkers function. P-values were adjusted based on Bonferroni correction and genes with at least 25% cluster-specific expression, >0.25 average log-fold change, and FDR<0.01 were chosen to identify the clusters. All cells were assigned scores based on cell type-specific gene modules to further identify broad cell types such as neurons, astrocytes, microglia and oligodendrocytes. A priori set of markers curated based on previous studies of cerebral organoids and developing fetal brain was explored to manually annotate the clusters. Additionally, we leveraged existing published fetal brain and organoid scRNA-seq datasets and undertook a correlation-based method^66^ (Spearman) to compare our annotated clusters to transcriptional profiles of previously annotated cell types.

### Differential expression testing and functional interpretation

We performed differential gene expression analyses on each cluster across conditions on log-normalized values using the MAST package^67^ in R. DEGs were considered significant if their Benjamini-Hochberg-adjusted P-value was < 0.05 and a log2 fold change was > 0.25. Significant DEGs were used to perform GO term overrepresentation analysis using the enrichR package. Gene set enrichment analysis (GSEA) was performed on ranked DEGs using fgsea package^68^ for pathway analysis (KEGG, GO:BPdatabase). Gene sets were limited by minSize=3 and nPerm=10000. Normalized enrichment scores were calculated and plotted for pathways with adjusted p-values < 0.05. Statistical significance of association of DEGs with gene signatures from published datasets was performed using Fischer’s exact test to obtain a p-value (<0.05, adjusted with BH method) and Odds ratio (OR). Number of genes expressed in the respective cell type was used as the genomic background.

### lntercellular interaction

Cellular crosstalk was inferred via ligand-receptor pair expression using CellphoneDB package^69^ (v2) in python. Statistical analysis was performed within the package with default parameters (1000 iterations) and no subsampling was done.

### Statistics

The assumptions of each used test were checked. All reported p-values are two sided and type of statistical test is reported in the figure legends or in the main text.

## DATA AND CODE AVAILABILITY

Processing of data and downstream analysis was performed in R (version 4.0.3). Figures used in this manuscript were generated in Python (version 3.6.12). Raw single-cell RNA sequencing data is deposited into GEO database (GSE181422). Source code to reproduce the findings are available in our repository on Github (https://github.com/SellgrenLab/organoid-Covid19). All other data is available from the corresponding authors upon request.

## ACKNOWLEDGEMENT

We thank the study participants. We are also grateful to the participant core facilities at Karolinska Institutet; The Biosafety Level (BSL)-3 laboratory at Biomedicum, BIC, and SciLife Lab. We are thankful to Judit Ozsvar, an intern at the Sellgren laboratory at time of the experiments, for her help with cell culture work, as well as to Gretchen Majkowitz for valuable feedback on the manuscript. This work was supported by grants from Hjärnfonden postdoktorala stipendier (S.: PS2019-0063), the Swedish Research Council (C.M.S.: 2017-02559), Karolinska Institutet (C.M.S.: KID), regional agreement on medical training and clinical research between Stockholm County Council (A.L.F, C.M.S.), One Mind Foundation/Kaiser Permanente (C.M.S.), and Marianne and Marcus Wallenberg Foundation (C.M.S.).

## AUTHOR CONTRIBUTIONS

C.M.S, S, A.O.O, S.K.G, and S.M conceived the project. R.H.P and S.D.S provided iPSCs. A.O.O derived the organoids. N.R.S and S performed infections, dissociations and related assays, supervised by A.R, with help from L.S. A.O.O, S, S.M optimized and performed IHCs, imaging and quantifications, with help from J.G.L. S.K.G optimized the qPCR assays and performed them with S.M. F.O derived microglia for 2D assays. S.M designed the scRNA-seq experiments with inputs from S and C.M.S. S.M analyzed scRNA-seq data. S and S.M interpreted the data. S, S.M, M.S, and C.M.S wrote the manuscript with inputs from the other co-authors.

