## supplementary for "SARS-CoV-2 promotes microglial synapse elimination in human brain organoids"

### Supplementary figure Legends

**Supplementary Figure 1.**

**(a)** 40X representative confocal image showing colocalization of astrocytic markers, GFAP and AQP4, in 110 DIV organoid. **(b)** qPCR analysis on supernatants of SARS-CoV-2 infected cerebral organoids at 56 DIV (n=3) showing log fold change in viral transcripts corresponding to nucleocapsid (N, black circle) and RNA-dependent RNA polymerase (Rdrp, turquoise square) genes. Also shown qPCR analysis on supernatants of infected VeroE6 cells for nucleocapsid (N, magenta diamond) gene. **(c)** NP-positive and Casp3-positive cells expressed as percentage of DAPI from two 56 DIV infected organoids (corresponding data points indicated by circles and triangles). Each data point is an average of 2-3 FOVs per section. Error bars indicate SEM. **(d)** Bar plot showing distribution of CASP3-positive cells amongst different cell-type markers, expressed as percentage of DAPI from two 56 DIV infected organoids. Mean represents an average of 4 FOVs representing different areas on the same section. Error bars indicate SEM. **(e)** 40X representative confocal images showing colocalization of CASP3 with neuronal- (MAP2 and PAX6) and glial- (GFAP and SOX10) lineage markers. **(f)** qRT-PCR on cell lysate fraction obtained following MACS with CD11b beads, showing expression levels of two microglial specific genes (*AIF1*, *TMEM119*). **(g)** Induced microglia (iMG) from a healthy donor, infected with SARS-CoV-2 and stained for dsRNA (for viral presence) and CASP3 (red), at 24hpi.

**Supplementary Figure 2. (a)** Scatter plot of percentage of mitochondrial reads versus number of expressed genes of the entire single-cell dataset prior to quality control. Red line indicated the applied cutoff for mitochondrial percentage. **(b)** Bar plot showing number of cells obtained per condition post quality control (see methods). Violin plots showing the distribution of transcript counts (left) and expressed genes per cell (right) across 6 samples on a **(c)** linear scale and **(d)** log scale [Control- Lane 1&2; 24hpi- Lane 3&4; 72hpi- Lane5&6]. **(e)** Violin plots showing the distribution of transcript counts (top) and expressed genes per cell (bottom) across every identified cell type cluster. **(g)** UMAP plots of overall embedding of the dataset integrated with two tools, CCA (left) and Harmony (right) as validation confirming no method-specific bias. Each cell is colored by its sample showing uniform distribution and no batch effects in the clustering analysis. **(h)** Bar plot with percentage cell type composition across individual samples. **(i)** UMAP plots displaying the distribution of QC metrics, percentage of mitochondrial reads (left), and calculated doublet score (right) per cell across all clusters.

**Supplementary Figure 3.** Cell type identification using a developmental single-cell dataset, *Pollen et al.,* 2019[1] (human primary and human organoids), as a reference dataset. Expression of top marker genes for each reference cell type (labelled on the left) are visualized across all cells on standard UMAP embeddings of our dataset. Cells are colored according to the estimated joint density of multiple marker genes represented by the scale on the right of each plot.

**Supplementary Figure 4. (a)** UMAP plots showing the expression of canonical markers genes across all cells. Spearman-ranked correlation of cerebral organoid clusters (rows) to reference transcriptomes (columns) of **(b)** human primary fetal cortex (*Nowakowski et al*., 2017),[3] **(c)** integrated human cerebral organoid datasets (*Tanaka et al.,* 2020), [2] **(d)** human primary developmental dataset (*Pollen et al.,* 2015).[5] **(e)** human primary developmental dataset (*Bhaduri et al.,* 2019),[4] **(f)** Dendrogram (left) and UMAP plot (right) integration space of microglial transcriptomic profiles comparing organoid-grown microglia to fetal (*Nowakowski et al*., 2017 & *Zhong et al., 2018*) and adult *(Hodge et al., 2019)* microglia. Cells are colored by their dataset. **(g)** UMAP plots highlighting a subset of neurons with *ACE2* expression along with expression of midbrain dopaminergic markers. This subset did not exhibit a differential response to SARS-CoV-2 in comparison to other neuronal clusters (Supplementary table S3). **(h)** IHC staining on 56 DIV organoid showing ACE2 expression (red) in B-III tubulin$^+$ neurons (green).

**Supplementary Figure 5. (a)** UMAP plot showing sub clustering analysis of microglial cells colored by their identified microglial subclusters and **(b)** condition (left), and sample (right). No differentially expressed genes (log2FC>0.2; adjusted p<0.05) were found between the microglial subclusters**. (c)** UMAP embedding of microglial subclusters displaying the distribution of QC metrics such as number of reads (nCount_RNA), number of expressed genes (nFeature_RNA), percentage of mitochondrial reads (percent.mito). **(d)** Heatmap of differentially expressed genes in microglia upon SARS-CoV-2 infection using MAST (see Methods). **(e)** Bar plots showing top significant pathways enriched for the two modules of genes found in microglia by over-representation analysis (Adjusted P<0.05). **(f)** Top significant upregulated and down-regulated KEGG pathways between SARS-CoV-2 exposed microglia (24hpi+72hpi) versus control microglia obtained by GSEA (adjusted P<0.05; see Methods). Color bar shows Normalized expression score (NES). Violin plots showing normalized expression of microglial markers associated with disease and homeostasis **(g)** and active phagocytic states **(h)** across conditions. **(i)** Dot plot displaying expression of canonical 'don't-eat-me' signals in neuronal clusters across infected conditions. (adjusted P<0.05). Size of the dot indicated fraction of cells expressing the gene. Scale bar indicated scaled-average expression values. **(j)** Quantification of IHC staining corresponding to Fig. 5d of $ISG15^+$ and $IBA1^+$ cells (Mann-Whitney U-test, P <0.0001).

**Supplementary Figure 6**. **(a)** UMAP embedding of astrocyte subclusters displaying the distribution of QC metrics such as number of reads (nCount_RNA), number of expressed genes (nFeature_RNA), and percentage of mitochondrial reads (percent.mito) per cell. **(b)** UMAP plot showing astrocyte subclusters with each cell colored by the sample. **(c)** Violin plots of cells scored by A1-reactive and A2-reactive signatures as defined in Barbar et al., 2020,[6] across astrocyte subclusters. **(d)** Astrocytic subclusters are visualized as columns on the heatmap with relative expression levels of top differentially expressed marker genes (rows) of individual subclusters (using MAST test implemented in Seurat, adjusted p-value<0.05). Top bar is colored by each astrocyte subcluster. High expression values shown in yellow while low expression values shown in purple. **(e)** Dotplot showing average-scaled expression of genes related to astrocyte activation and function across astrocyte subclusters.

**Supplementary Figure 7. (a)** UMAP embedding of choroid plexus subclusters displaying the distribution of QC metrics such as number of reads (nCount_RNA), number of expressed genes (nFeature_RNA), percentage of mitochondrial reads (percent.mito), and calculated G2M per cell. **(b)** UMAP plot showing choroid plexus subclusters with each cell colored by the sample. **(c)** Heatmap showing scaled expression of top differentially expressed markers by choroid plexus subclusters (obtained using MAST test implemented in Seurat, adjusted p-value<0.05). Top bar colored by each choroid plexus subcluster. High expression values shown in yellow while low expression values shown in purple.
